# Plastimos: a live cell imaging-based framework to study dynamics of EMT-mediated cellular plasticity in breast cancer

**DOI:** 10.1101/2025.05.28.656591

**Authors:** Saba Sameri, Durdam Das, Silvia Materna-Reichelt, Nataša Stojanović Gužvić, Lukas Wöhrl, Martin Hoffmann, Hedayatollah Hosseini

## Abstract

Cancer cell plasticity, primarily mediated by the process of epithelial-mesenchymal transition (EMT), plays a critical role in promoting therapeutic resistance, tumor heterogeneity, and metastasis. EMT-mediated plasticity enables cancer cells to undergo molecular and phenotypic changes, which alter their invasiveness and resistance to treatments. This study aims to develop a quantitative, high-throughput system to assess EMT-mediated plasticity, which can inform therapeutic strategies for metastatic cancers. To accomplish this, we developed Plastimos, a semi-automated imaging analysis pipeline. This framework utilizes time-series images to track live cells through deep learning-based segmentation and employs a greedy algorithm to map cell trajectories, enabling the extraction of cellular phenotypic features. These features are used to study the EMT-mediated plasticity state of cells in response to the well-known EMT-inducing factors EGF and TGF-β1. We selected two breast cancer cell lines, MCF7 and MDA-MB-231, representing classical epithelial-like and mesenchymal-like cell. The pipeline assigns a Plasticity Index based on various parameters, including motility, morphology, and proliferation, thus providing a quantitative estimate for the plasticity of deviating from the epithelial state. Our results indicate that epithelial-like cells respond to EMT-inducing factors at both molecular and phenotypic levels, while mesenchymal-like cells’ responses are only seen phenotypically. The Plasticity Index classifies cells along the EMT spectrum, converting to a Plasticity Score that reflects mesenchymal proportions.

**Availability and implementation:** The pipeline implementation and the source code of Plastimos can be found at: https://github.com/Durdam/Plastimos.git

## Introduction

Cancer cell plasticity is defined by the ability of malignant cells to change and adapt their morphology and function in response to environmental factors within the tumor microenvironment [1]. This property is critical to understand tumor heterogeneity, metastasis, and treatment resistance, all of which are fundamental to the clinical course of cancer [1–3].

Epithelial-mesenchymal transition (EMT) is widely known as the major driver of cancer cell plasticity. EMT is a biological process wherein epithelial cells lose their cell-to-cell adhesion properties along with transcriptional reprogramming, adopting a mesenchymal phenotype that enhances motility and invasiveness [4, 5]. During EMT, certain transcription factors become activated, promoting the aggressiveness and stemness of cancer cells [2, 4].

The characterization of cancer cell plasticity, especially within the context of EMT, is essential for developing effective anti-metastatic therapies [8, 9]. Live-cell imaging offers a unique advantage by allowing real-time monitoring of cellular dynamics, capturing how cells interact and change in response to distinct stimuli [10]. As cells transition through EMT, they exhibit notable morphological changes, such as alterations in shape and size, which traditional microscopy methods may struggle to capture [11, 12]. Moreover, EMT is a spectrum rather than a binary process, and intermediate states of plasticity are still not fully understood. Since mortality in cancer patients typically results not from the primary tumor (PT) itself, but from metastatic spread (mets), effective drug screening protocols should extend beyond terminal viability assessments to incorporate real-time quantification of metastatic potential, highlighting the need for further research of EMT’s states and their clinical relevance [6].

Image-based machine learning (ML) approaches, which are developing rapidly with the emergence of widespread artificial intelligence (AI) applications, are powerful tools for processing large datasets. New ML-based image analyses surpass traditional methods in predictive accuracy. By incorporating multiple inputs, ML models improve our understanding of EMT and its associated cellular changes [13]. The field of automated cell tracking has contributed valuable insights into various cellular dynamics. The emergence of new methods that improved the resolution, dimensionality, extent and throughput of optical microscopes demands new, improved tracking algorithms. Furthermore, the rapid evolution of ML [14] is changing the way cell tracking is performed, as deep neural networks swiftly replace classical image analysis methods. These models provide impressive results while posing their own share of challenges related to training strategies, quality and quantity of available data, parametrization, and generalization [15]. The development of deep learning (DL) holds great promise to further improve predictive modeling in oncology. This advancement may facilitate more precise treatment and, as a result, improved patient care [16–18].

In this study, we developed an imaging-based pipeline for monitoring the *in vitro* plasticity of cancer cells, utilizing DL-based segmentation and live-cell tracking features implemented in R. The segmented masks are further used as input to a simple yet precise greedy algorithm to track cells in time series imaging data and extract single cell features. These features include morphology, motility, and growth rate and represent the cells along the EMT spectrum. Instead of categorising cells into discrete classes of EMT states [19], we used these features to define a continuous spectrum of EMT. We incorporated EGF (epidermal growth factor) and TGF-β1 (transforming growth factor beta1) stimulation, as two well-known EMT-inducing factors [20– 24]. Our assay models EMT-associated plasticity by comparing highly metastatic mesenchymal-like cells to low-metastatic epithelial-like counterparts. The pipeline outputs a Plasticity Index (PI), quantifying metastatic potential *in vitro*. Averaging PI values across single cells generated a plasticity score per cell line and treatment. Dimensionality reduction of PI features confirmed the method’s capacity to resolve phenotypic differences between cell types. This strategy enhances understanding of cancer cell plasticity and supports *in vitro* evaluation of anti-metastatic therapies.

## Methods and Material

### Cell Culture and Reagents

Human breast cancer cell lines MCF7 and MDA-MB-231 were obtained from ATCC and cultured in monolayer as recommended by the provider. MCF7 cells were grown in RPMI 1640 medium (Anprotec; AC-LM-0060) with 10% FBS (Pan-Biotech; P40-37500), 2 mM L-glutamine (Pan-Biotech: P04-80100), and 10 unit/mL penicillin/streptomycin (Pan-Biotech: P06-07050). MDA-MB-231 cells were maintained in DMEM High Glucose medium (Anprotec; AC-LM-0011) supplemented similarly. Cell line authenticity was confirmed via STR analysis (Cell-ID™, Promega). Cultures were kept at 37°C, 5% CO2, and routinely tested negative for mycoplasma. TGF-β1 1 (HEK293 derived) (Peprotech; 100-21) and hEGF (Sigma-Aldrich; E9644) were prepared as per manufacturer’s instructions and replenished daily.

### Generation and propagation of breast cancer tumoroid models in suspension culture

Breast cancer cell lines MCF7 (RPMI, 10% FBS, 1 x P/S, 1x Gl-Max, 0.01 mg/ml insulin) and MDA-MB-231-GFP (DMEM, 10% FBS, 1 x P/S, 1x Gl-Max, 1.5 µg/ml puromycin) were grown in 2D cultures in standard T25 cell culture flasks under standard conditions (5% CO2, 37°C). To start the 3D culture, single cells were obtained by Trypsin/EDTA treatment of the ~ 80% confluent monolayer.

Low-viscosity matrix (LVM) for 3D suspension culture was prepared by mixing breast cancer tumoroid medium [25] with 5% v/v Geltrex (Geltrex LDEV free RGF BME, Thermo Fisher Scientific; A1413202) on ice. Single cells were mixed with the medium and plated onto ultra-low attachment coated multiple well cell culture plates (Costar 6-well; 3471 or Costar 24-well; 3473) based on a 3D suspension culture protocol [26]. For 24-well plates, 25000 single cells were plated in 500 µl medium and 200000 single cells for 6-well plates in 2 ml medium. Two times per week, 250 µl/ 1 ml medium was added per well in 24-/ 6-well plate and mixed by pipetting up and down several times.

During the time of culturing, brightfield images were obtained to monitor the formation of tumoroids (Olympus IX81F-3).

For passaging, the formed tumoroids from 24- or 6-well plates were transferred into 2 ml or 5 ml tubes on ice. The tubes were centrifuged for 5 minutes at room temperature, 300 x g. The supernatant was removed and 200 µl or 1000 µl TrypLE (ThemorFisher Scientific; 12605028) was added for ~ 10-15 minutes, incubated at 37°C, 5% CO2 and mixed every 5 minutes. After dissociation into single cells, 1 ml or 4 ml ice cold advanced DMEM/F12 (1x P/S, 1 mM Hepes, 1x Gl-Max) was added and cells were counted and prepared for passaging.

### RNA Analysis

Total RNA was isolated using the NucleoSpin RNA Mini Kit (Macherey-Nagel:740955.50) according to the manufacturer’s protocol. Complementary DNA (cDNA) was synthesized using a reverse transcriptase kit (ThermoFisher: k1622). Quantitative PCR was performed on a LightCycler 480 instrument (Roche) using Fast Start Master SYBR Green Kits (Roche; 4707516001), with 5 ng of cDNA per reaction. Human reference cDNA served as a positive control. Measurements showing unspecific products in melting curve analysis were excluded. Gene expression levels were normalized to β-actin (Actb). Relative expression was calculated using the 2^−ΔΔCt^ method [27]. All primers were obtained from Eurofins MWG Operon, Germany and all primer sequences are provided in Table 2.

### Immunofluorescence (IF) Staining

For monolayer cultures, cells were seeded in 24-well plates at 4 × 10^3^ cells per well. After 72 hours (including 24 hours post-seeding without treatment and 48 hours with treatment), cells were washed with PBS 1X, fixed with 4% paraformaldehyde for 10 minutes, and washed again. Permeabilization was performed using 0.3% Triton X-100 (Sigma: T8787) in TBS, followed by washing with TBS 0.1% Triton X-100 and blocking with Goat Serum (DAKO) in TBS 0.1% Triton X-100 at room temperature. Primary antibody incubation (details in Supplementary Table 3) was conducted for 1 hour at room temperature. After washing with TBS 0.1% Triton X-100 four times, cells were incubated with fluorescently labeled secondary antibodies (Invitrogen) and Hoechst 3342 (1:1000, ThermoFisher: 62249) for 1 hour at room temperature. The wells were washed then three time with TBS 0.1% Triton X-100 and one time with PBS 1X. The images were acquired using Olympus IX81.

### Pipeline for Sphere quantification

We obtained tumoroid culture data for various time points, using Olympus IX81F-3, starting from day 3 to day 8 for both MCF7 and MDA-MB-231 cell line models. To analyze these images, we first created a segmentation mask using the otsu threshold-based algorithm. However, the masks were not very accurate. For this reason, we used the Cellpose GUI (Graphical User Interface) to remove background debris and improve the cell masks. Since, we wanted to assess the protrusionlike structures of the tumoroid, it was crucial that the masks accurately captured the branching structures next to the main cell body. After the segmentation, the masks were used as input to the “MeasureObjectSkeleton” module in cellprofiler [28] to quantify the number of protrusions coming out of the cells, which were then compared between MCF7 and MDA-MB-231 untreated cells.

### Cell motility assay

We performed time-lapse live cell imaging using Incucyte® S3 Live-Cell Analysis System (Sartorius) allowing monitoring the migration behavior of different cells in real time. To this end, MCF7 and MDA-MB-231 cells were cultured in 6-well plates (30,000-35,000 cells/well) and treated with either 5 ng/mL of TGF-β1 or 100 ng/mL of EGF for 24 h. Subsequently, the medium was exchanged with fresh medium containing the treatments. The plate was placed in the Incucyte for 26 h at 37°C and 5% CO_2_ and imaged with brightfield every 10 minutes totaling 150 frames of imaging data for each of 3-4 replicates taken for both cell lines. The number of replicates varies between treatment conditions as in some replicates the images were not of sufficient quality.

### Cell tracking pipeline

Images were first segmented by Cellpose using the cyto3 model and further fine-tuned by retraining on a small subset of manually labelled masks. This helped generating accurate masks for the rest of the images. Post segmentation, the masks were fed into the cell tracking pipeline that uses a greedy algorithm to map the trajectory of the cells.

First, each cell in a frame is assigned a unique integer label. We then used the Jaccard index to measure the similarity to each cell in the consecutive frame in the forward direction. In case, there are multiple cells overlapping in the next frame, we take the cell with the highest similarity to map the path. This method worked well for our image data, since the cells were seeded in low density. In case the cells were in very high density, there would be chances that we encounter some errors in tracking. The time gap between frames is also crucial in this respect. If a cell travels a large distance from one frame to another resulting in no overlap between frames, the path might be lost. Therefore, by taking images every 10 minutes, we made sure that there is always some overlap across frames. This also helped to eliminate debris and artifacts since these move quite fast resulting in no overlap across frames and thus get eliminated in the tracking pipeline that requires an object to persist for at least seven consecutive frames. The Incucyte camera captures the center field of the plate Thus, requiring seven consecutive frames also largely protects against reentry events. Seven is an arbitrary number that we decided for after intensive cross checking our imaging results. To monitor morphological cell features across frames we used functions from the EBImage R-package [29]. Further details and the implementation of the pipeline is available on the Github repository.

### Statistical Analysis

Statistical analyses within each dataset were conducted using GraphPad Prism V9.5. All experiments were independently performed at least three times. To determine whether parametric or non-parametric tests were appropriate, D’Agostino normality tests were conducted if the number of data points was sufficient for testing. For cases with a low number of data points, where D’Agostino test could not be performed, the Shapiro-Wilk test was applied to assess the normality distribution of the sample. To compare changes in mean values between treated groups with the control groups, one sample t-tests were utilized for data that passed the normality test, and the Wilcoxon tests were used otherwise. For multiple comparisons, Kruskal-Wallis tests were employed. Spearman correlation tests were used to evaluate correlations between traits. All p-values presented in the figures and legends were calculated two-tailed.

## Results

### Overview of the pipeline for analyzing the EMT-mediated plasticity of cancer cells

To establish a baseline for evaluating the plasticity state of cancer cells and its correlation with metastatic potential, breast cancer was selected due to its heterogeneity a and presence of multiple molecular subtypes, each with unique biological and clinical behavior [30](Fig. 1A). Our strategy was to select representative *in vitro* models of two breast cancer subtypes with significant differences in metastatic potential. The cells were tracked for 25 hours in culture plates and exposed to two EMT inducers generating dynamic shifts. To prevent restrictions due to high cell density, cells were cultured at a low density, allowing ample space for cellular movement and morphological changes. Cells were imaged in brightfield microscopy every 10 minutes during the observation period (Fig. 1A).

**Figure 1.**
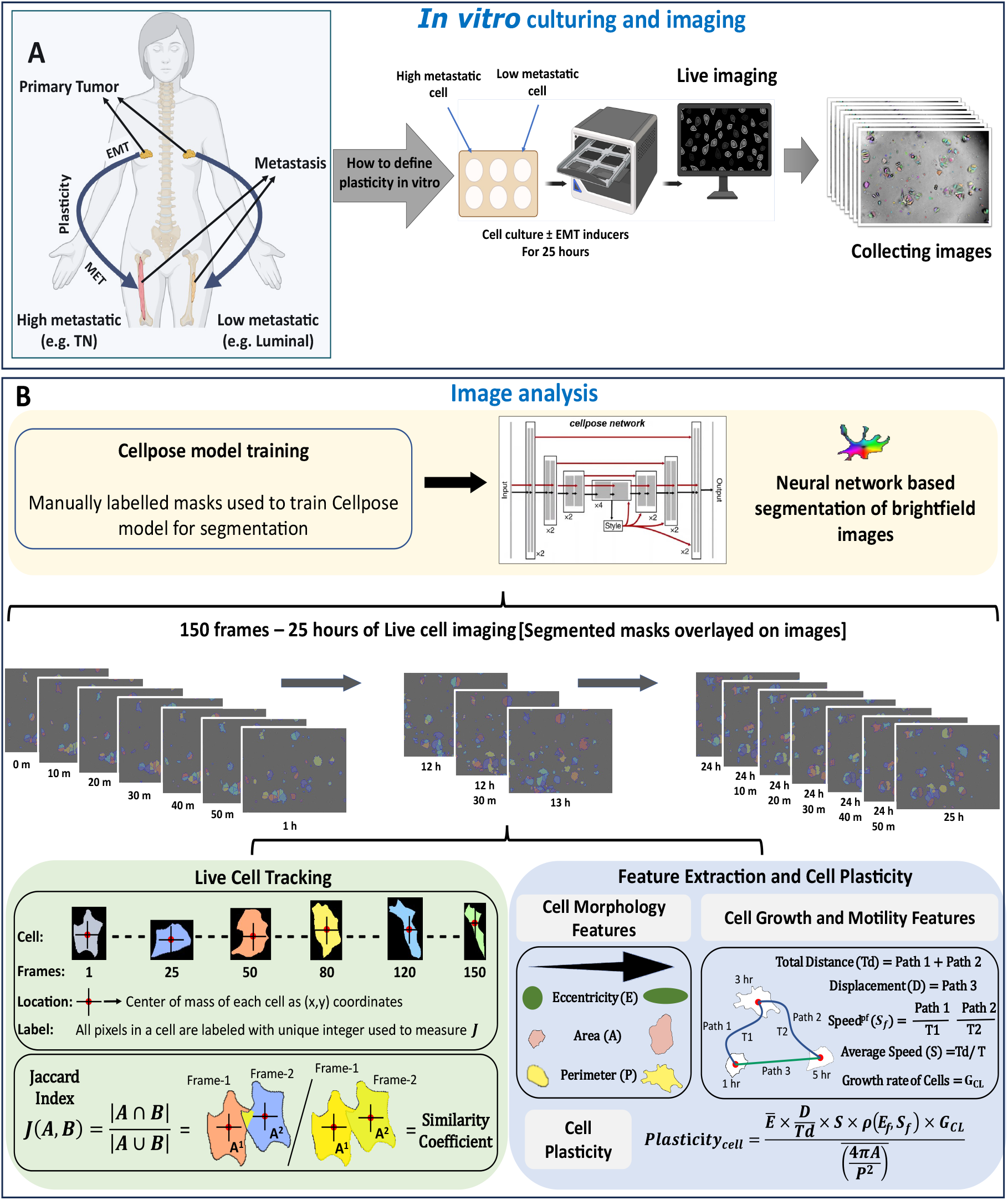
Overview of the experimental approach for quantifying EMT-mediated cancer cell plasticity through integrated live cell imaging and computational analysis. **A**) Schematic representation of cancer cell plasticity and metastasis progression in breast cancer. Metastasis in patients depends on the EMT-mediated plasticity program of cancer cells, which involves a dynamic transition between EMT in the primary site and its reversal process, MET, at metastatic sites. Two distinct breast cancer phenotypes are highlighted: highly metastatic (e.g., triple-negative) and less metastatic (e.g., luminal) subtypes, demonstrating differences in plasticity and metastatic potential, followed by bright-field live-cell imaging (right panel). **B**) Computational pipeline for analyzing cancer cell plasticity from *in vitro* models. The upper panel shows the workflow from manual mask labeling to neural network-based segmentation using the Cellpose framework. The middle panel displays the temporal sequence of 150 frames captured over 25 hours of imaging, with segmented mask overlays demonstrating successful automated cell detection. The lower left panel illustrates the live cell tracking methodology, where individual cells are tracked across 150 frames with the Jaccard Index measuring cell similarity. Cells are assigned unique labels, and their location is determined by the (x,y) coordinates of their center of mass. The lower right panel details the feature extraction process, presenting the quantification of cell morphology features (Eccentricity, Area, Perimeter), cell growth (Growth rate,) and motility features (Total Distance, Displacement, Average Speed). The final equation at the bottom right integrates these parameters into a comprehensive scoring metric called Plasticity Index, incorporating both morphological and behavioral metrics to quantify cellular plasticity *in vitro*.

As a measure of the plasticity state of cells, we propose that plasticity can be characterized by motility, morphology, and proliferation characteristics. The morphology features were cell eccentricity (E), compactness, area (A), and perimeter (P) since alterations in these features are known to capture metastatic behaviour [32]. For assessment of cell motility, that is often related to EMT changes [33], we tracked the total distance (Td), the overall displacement (D), and the speed per frame (S_f_) and on average (S). Cell proliferation (G_CL_) was also quantified. Finally, the above parameters were combined in an equation to calculate an overall PI for each cell (equation 2) (Fig. 1B, lower right panel).

### Selecting and validating cell line models

The MDA-MB-231 and MCF7 cell lines are commonly used to represent breast cancer subtypes with different levels of metastasis potential. The highly metastatic triple-negative subtype MDA-MB-231 and the low metastatic luminal subtype MCF7 differ in morphology and behaviour in ways that render them excellent models for the development and validation of our imaging-based analysis pipeline [4, 32, 34–38]. MCF7 cells display epithelial characteristics, such as cell clustering and cuboidal shapes, while MDA-MB-231 cells exhibit mesenchymal traits, including an elongated, spindle-like appearance and lack of cohesive cell clustering [4]. Based on previous studies, we developed a standardized metastatic potential scale that ranks multiple breast cancer cell lines according to their documented invasive behaviors (Fig. 2A). This classification framework distinguishes between non-metastatic, poorly metastatic, intermediately metastatic, and highly metastatic phenotypes across the breast cancer cell line spectrum. This analysis confirmed our expectations and helped to solidify MDA-MB-231 cells as highly metastatic and MCF7 cells as poorly metastatic as ideal comparative models for our subsequent analyses [4, 33, 36–38].

**Figure 2.**
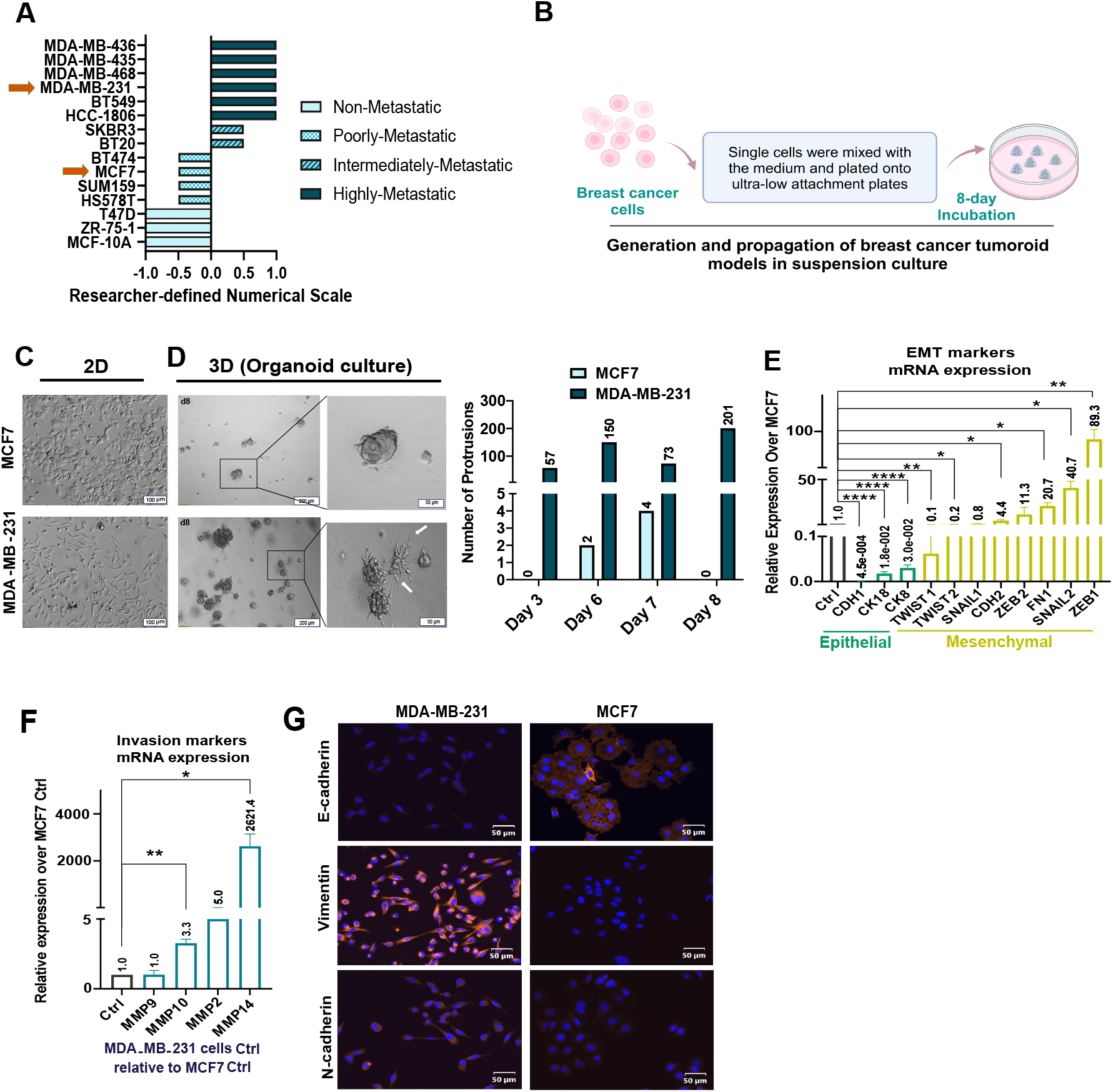
Characterization of morphological and molecular features distinguishing MCF7 and MDA-MB-231 breast cancer cell lines. **A)** Comparative analysis of metastatic potential across multiple breast cancer cell lines (1), classified into four categories: non-metastatic, poorly metastatic, intermediately metastatic, and highly metastatic based on a researcher-defined numerical scale (−1.0 to 1.0) that quantifies their relative metastatic capacity. MDA-MB-231, as a highly metastatic (Index +1.00), and MCF7, as a poorly metastatic (Index −0.5) cell line were selected for this study. **B)** Schematic representation of the experimental workflow for generating and propagating breast cancer cells in suspension culture. Single cells were mixed with the tumoroid medium and plated onto ultra-low attachment coated multiple well cell culture plates for 8 days. During the time of culturing, bright field images were obtained to monitor the formation of tumoroids (See Method section). **C)** The cellular morphology of MCF7 (cuboidal) and MDA-MB-231 (fibroblast-like) cells in 2D. Scale bars = 100 µm. **D)** Protrusions in 3D culture of MDA-MB-231 and MCF7 cells. Quantification of cellular protrusions in both cell lines across different time points (D, right panel). Scale bars = 200 and 50 µm. **E)** The mRNA expression (qPCR results) of EMT (epithelial and mesenchymal) markers and **F)** invasion markers in MDA-MB-231 cells (no treatment), relatively quantified to MCF7 cells (no treatment). **G)** Immunofluorescence imaging showing differential expression of key EMT markers (E-cadherin, Vimentin, and N-cadherin) in MCF7 and MDA-MB-231 cells. Blue = DAPI (nuclei), Red = target proteins. Scale bars = 50 µm. Statistical significance in panels E and F was assessed using One Sample t-tests. Significance levels are indicated as follows: P > 0.05; *: P ≤ 0.05; **: P ≤ 0.01; ***: P ≤ 0.001; ****: P ≤ 0.0001.

To better define the differences, we assessed the cellular characteristics of MCF7 and MDA-MB-231 cells grown in both 2D and 3D culture (Fig 2B-D). In 2D culture, MCF7 cells exhibited a typical epithelial morphology with a cobblestone-like arrangement, whereas MDA-MB-231 cells displayed a mesenchymal, elongated phenotype. These discrepancies were even more evident in 3D tumoroid culture where MDA-MB-231 cells displayed a higher number of protrusions than MCF7 cells over the time course (Days 3-8). Quantitative analysis showed that MDA-MB-231 cells attained up to 200 protrusions on Days 6 and 8, whereas MCF7 cells showed sparse protrusion formation (Fig. 2D, right panel).

Molecular profiling using qPCR further highlighted these phenotypic distinctions, especially in the mRNA expression of EMT markers (Fig. 2E). The mRNA expression analysis demonstrated significantly elevated levels of mesenchymal markers in MDA-MB-231 cells compared to MCF7 cells. Similarly, analysis of invasion marker’s mRNA expression revealed a strong upregulation in MDA-MB-231 cells, with some markers being over 2000-fold more expressed than in MCF7 cells (Fig. 2F). Immunofluorescence analysis confirmed these observations, as protein expression patterns of classical EMT markers distinctly differed between the two cell lines (Fig. 2G). MCF7 cells expressed high levels of E-cadherin protein (epithelial marker) and low levels of vimentin and N-cadherin proteins (mesenchymal markers) making the cells distinctly epithelial. MDA-MB-231 cells, on the other hand, displayed strong vimentin expression and low E-cadherin, consistent with a classical mesenchymal phenotype.

Collectively, these findings show that MCF7 and MDA-MB-231 cells are located at opposite ends of the EMT spectrum. MDA-MB-231 cells are aggressive, mesenchymal, while MCF7 cells are non-aggressive, epithelial. This separation of both, morphology and molecular signatures, confirms the eligibility of chosen model system to study cellular plasticity in the context of EMT in breast cancer.

### TGF-β1 and EGF differentially regulate EMT programs in MCF7 and MDA-MB-231 cells

We hypothesized that treating our cell line models with classical EMT regulators will enhance the ability to more precisely and digitally track and record changes in the plasticity potential of the tested cells. To prove this, we treated MCF7 and MDA-MB-231 cells with TGF-β1 and EGF as two well-known EMT-inducing factors [39, 40]. Immunofluorescence analysis revealed distinct patterns of EMT marker expression between the two cell lines under these treatment conditions. In MCF7 cells, TGF-β1 treatment induced a partial EMT response characterized by decreased E-cadherin expression and increased N-cadherin levels. EGF treatment did not significantly affect EMT markers in MCF7 cells (Fig. 3A). MDA-MB-231 showed no noticeable changes in the protein levels of EMT markers under TGF-β1 and EGF treatments (Fig. 3B).Quantitative PCR analysis of EMT markers in cells treated with TGF-β1 for 48 h revealed significant changes in mRNA expression levels, compared to the untreated control cells (Fig. 3C-D). MCF7 cells showed a marked shift in their EMT profile upon TGF-β1 treatment, with upregulation of mesenchymal markers’ mRNA levels and no significant change in epithelial markers. MDA-MB-231 cells maintained their mesenchymal profile under TGF-β1 treatment. Analysis of invasion markers demonstrated that TGF-β1 treatment for 48 h significantly increased the mRNA expression of invasion-associated genes in MCF7 cells but had no significant effects on MDA-MB-231 cells (Fig. 3E-F). Interestingly, MDA-MB-231 cells, which already exhibited a mesenchymal phenotype, showed no further enhancement of mesenchymal markers at the protein and mRNA levels under both TGF-β1 and EGF stimulation, which may reflect a saturation level of mesenchymal regulatory mechanisms in this highly mesenchymal state cell line. To explore this, we additionally quantified the mRNA expression of EMT and invasion markers in MDA-MB-231 cells treated with TGF-β1 relative to MCF7 cells also treated with TGF-β. These results showed that, although the response of MCF7 cells to TGF-β1 was more pronounced than that of MDA-MB-231 cells, MDA-MB-231 still displayed more mesenchymal properties compared to the MCF7 cell line (Sup. Fig. 1A-B).

**Figure 3.**
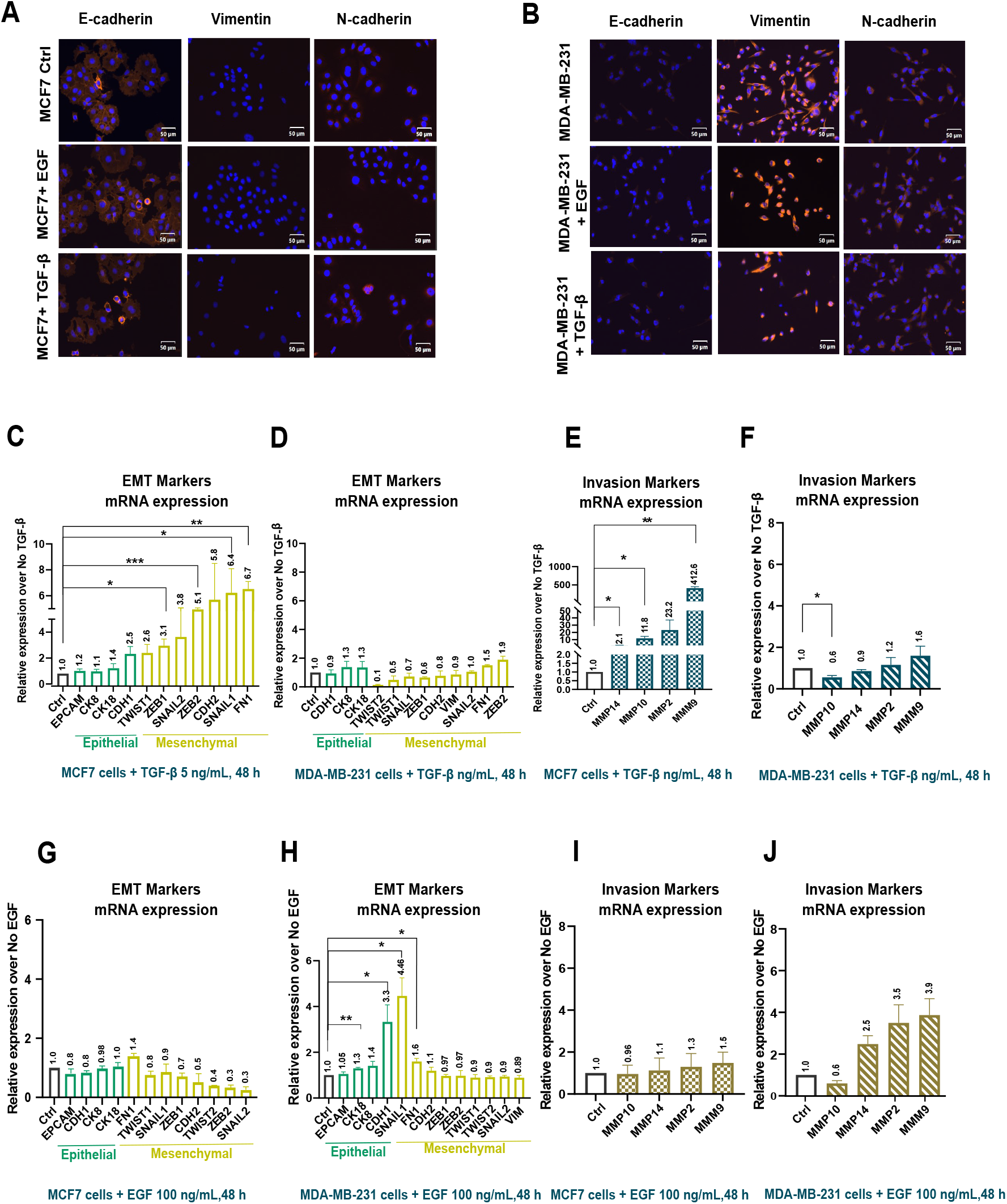
Differential EMT responses to TGF-β and EGF treatment. **A-B**) Immunofluorescence imaging of EMT markers in MCF7 cells (A) and MDA-MB-231 cells (B) under control conditions and following treatment with EGF or TGF-β. Expression patterns of E-cadherin, Vimentin, and N-cadherin are shown. Blue = DAPI (nuclei), Red = target proteins. Scale bars = 50 µm. **C-F**) Quantitative mRNA analysis (qPCR) of EMT and migration/invasion markers’ expression in response to TGF-β treatment. Relative expression levels of epithelial and mesenchymal markers in MCF7 cells (C) and MDA-MB-231 cells (D). Relative expression levels of migration/invasion markers in MCF7 cells (E) and MDA-MB-231 cells (F). **G-J**) Quantitative mRNA analysis (qPCR) of EMT and migration/invasion markers’ expression in response to EGF treatment. Relative expression levels of epithelial and mesenchymal markers in MCF7 cells (G) and MDA-MB-231 cells (H). Relative expression levels of migration/invasion markers in MCF7 cells (I) and MDA-MB-231 cells (J). Statistical significance in all panels was determined using One Sample t-tests for panels C,E,F,H,I and J and Wilcoxon tests for panels D, and G. Significance levels are indicated as follows: P > 0.05; *: P ≤ 0.05; **: P ≤ 0.01; ***: P ≤ 0.001; ****: P ≤ 0.0001.

The response to EGF treatment showed distinct patterns compared to TGF-β1 stimulation stimulation (Fig. 3G-H). In MCF7 cells treated with EGF the data reveal a notable decrease in EMT marker expression, although statistical analysis did not reach significance (Fig. 3G). Mesenchymal markers (with the exception of TWIST1 and SNAIL1 and FN1) consistently show downregulation compared to control. Meanwhile, epithelial markers maintain relatively stable expression levels (Fig. 3G). This pattern suggests that EGF treatment may preferentially suppress mesenchymal characteristics while preserving epithelial marker expression in MCF7 cells. MDA-MB-231 cells showed enhanced expression of certain mesenchymal markers (SNAIL1 and FN1) and epithelial markers (E-cadherin (CDH1) and CK18), under EGF treatment over 48 h (Fig. 3H). The invasion marker profile under EGF treatment revealed no significant changes in MCF7 cells and MDA-MB-231 cells (Fig. 3I-J). We then quantified the mRNA expression of EMT and invasion markers in MDA-MB-231 cells treated with EGF relative to MCF7 cells also treated with EGF. Consistent with the results for TGF-β1 (Sup. Fig. 1A-B), MDA-MB-231 cells treated with EGF exhibited more mesenchymal and invasive features compared to MCF7 cells (Sup. Fig. 1C-D).

These findings show that MCF7 and MDA-MB-231 cells have different sensitivities to EMT-inducing factors. Our findings indicate that TGF-β1 (5 ng/mL) generally provokes a stronger EMT response than EGF (100 ng/mL) in both epithelial-state and mesenchymal-state cells. Specifically, our findings indicate that EGF functions as an EMT inducer in mesenchymal-state cells but as an EMT inhibitor in epithelial cells. Differential responses of these cell lines to the treatments further support our method to investigate alterations of the plasticity program in our cancer models.

### Quantitative analysis of plasticity index features

Our preliminary results facilitated the selection of study models with the appropriate EMT phenotypic and genotypic characteristics and their alteration under experimental treatments. This ensured alignment with our objectives and allowed to accurately assess the plasticity potential of cancer cells *in vitro*. We performed comprehensive analyses of the live cell imaging data and extracted features including motility, proliferation and cell morphology. Cell motility features like total distance covered by a cell, displacement, average speed and combined features like displacement-to-distance ratio of a cells (Fig. 4A-D) distinctly capture the difference between high epithelial-like cells of the MCF7 population and high mesenchymal-like cells of the MDA-MB-231 population. Cells show a high migratory effect with the treatment of EGF and TGF-ß compared to the untreated cells in both MCF7 and MDA-MB-231 cell lines. The highest difference between the six conditions is found in the average speed of cells shown in figure 4D as all the conditions are significantly different from one another. In both cell lines, the treatment of EGF and TGF-ß enhance the cell movement. The speed was measured per frame as well (Sup. Fig. 2A) and shows a similar pattern to the average speed.

**Figure 4.**
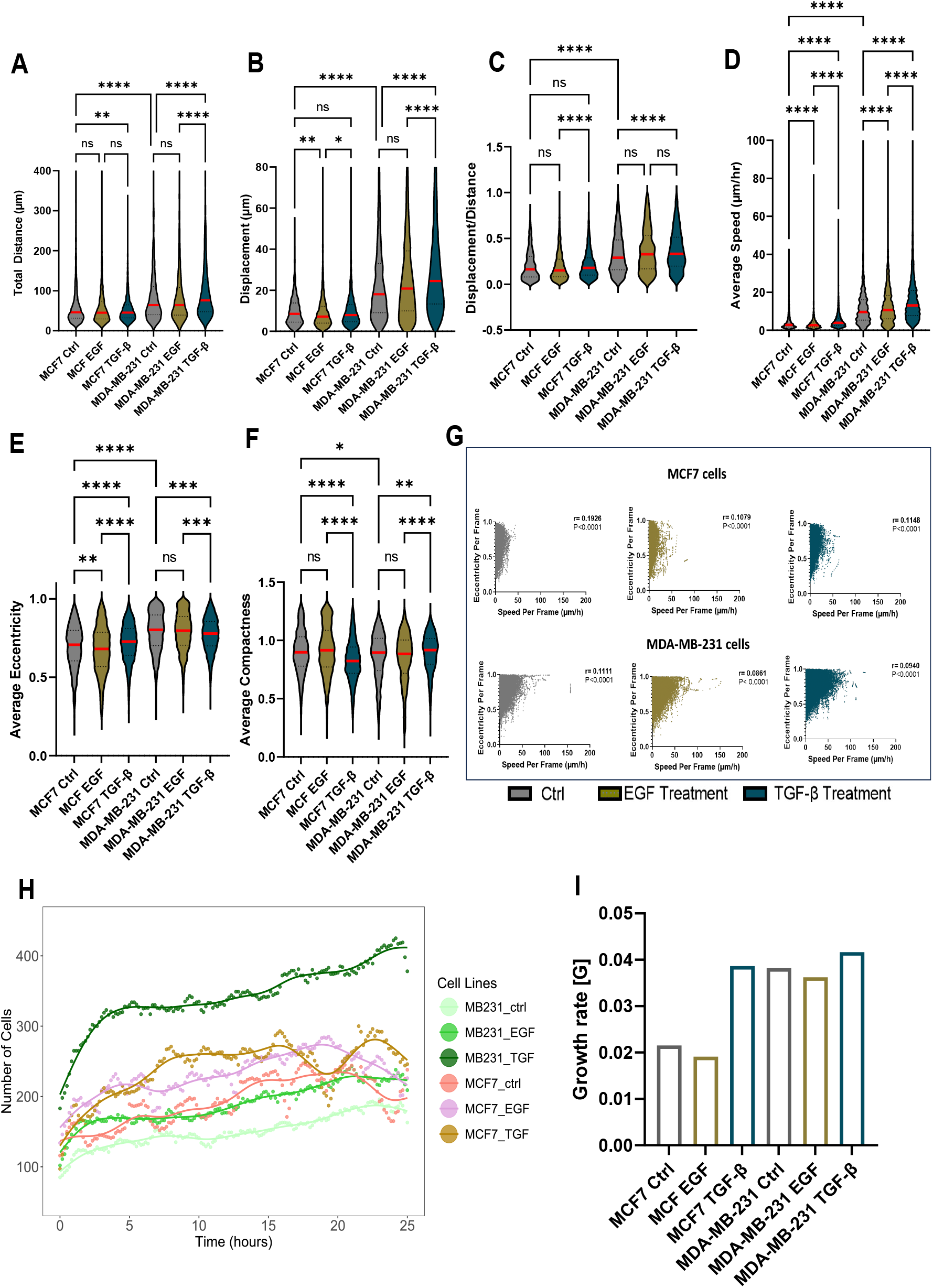
Quantitative analysis of cell motility and morphological features under EGF and TGF-β treatment. **A**) Total distance traveled by cells under different treatment conditions, showing comparative analysis between MCF7 and MDA-MB-231 cells in control conditions and following EGF or TGF-β treatment. **B**) Cell displacement measurements across all experimental conditions, quantifying the straight-line distance between start and end points of cell movement. **C**) Displacement to Total Distance ratio. **D**) Average speed measurement of each cell across all frames they were tracked through. E) Eccentricity measurements quantifying cell shape changes. F) Analysis of average compactness of each cell. G) Relation between Eccentricity per frame and Speed per frame features shown as scatter plot with spearman correlation and p-value indicated on the top right of all the subplots. H) Number of cells detected at each time point up to 25 hours show as dots. Fitted curves using smoothing splines for each condition are shown as lines. I) Average growth rate G measured for all conditions using the fits from H) shown as bar-plot. All violin plots show data distribution with median values indicated. All data are presented as violin plots showing data distribution, with median values indicated. Comparisons were made between treatment groups and across cell lines using Kruskal-Wallis tests. Significance levels are indicated as follows: ns (not significant): P > 0.05; *: P ≤ 0.05; **: P ≤ 0.01; ***: P ≤ 0.001; ****: P ≤ 0.0001.

The morphological features include eccentricity, which measures how much an object’s shape deviates from being spherical, and compactness, which measures how closely packed a shape’s area is. Eccentricity of a cell was calculated in each frame (Sup. Fig. 2B) and the average eccentricity is shown in figure 4E for all the six conditions. Interestingly, the EGF treatment seems to reduce the median eccentricity for both MCF7 and MDA-MB-231 cells. For the MDA-MB-231 cells, which are already in a mesenchymal state, the treatment of EGF and TGF-ß reduces the median average eccentricity indicating that these cells are not getting further elongated. The compactness was also measured in each frame (Sup. Fig. 2E) and further the average compactness was calculated (Fig. 4F). MCF7 with TGF-ß treatment has the lowest median average compactness across all six conditions indicating higher irregular shapes with filopodia like structure in the cell membrane as compared to more smooth and round shaped membrane found in the MCF7 control condition. However, the MDA-MB-231 only show minor changes in the compactness between the control and the two treatments. Further, area and perimeter features of the cells were also extracted, (Sup. Fig. 2C-D) and both these features show a similar pattern where the MDA-MB-231 cells are generally larger than the MCF7 cells. Interestingly, EGF and TGF-ß treatments slightly increase median cell size in MDA-MB-231 cells but decrease it in MCF7 cells, compared to their respective controls as shown in the area and perimeter features.

These results are consistent with our gene expression analyses, where we observed that both EGF and TGF-β1 stimulate the upregulation of mesenchymal and/or invasion markers in MDA-MB-231 cells, whereas this induction was observed mainly with TGF-β1 in MCF7 cells (Fig. 3C-J and Sup. Fig 1 A-C).

Since EGF and TGF-ß affects both morphology and cell speed, it was intriguing to determine whether these features have any correlation. As stated by (Quinsgaard et al., 2024), both speed and morphology change in response to EGF and TGF-ß treatments but are not dependent on one another. Contrary to that, we do find a very interesting pattern when we compared a morphological feature like eccentricity and a motility feature like speed (Fig. 4G). For the high epithelial-like cells of MCF7 control as well as EGF and TGF-ß treatment most cells have a maximum speed up to 50 μm/h, whereas in all MDA-MB-231 categories, the maximum speed goes up to 100μm/h and we see a clear upside-down right triangle shape for all the three MDA-MB-231 conditions in the scatterplot. Although the overall correlation is not very significant, there seems to be some correlation between the cell’s morphology and movement for high mesenchymal like cells. These results suggest that the type of morphological changes may vary for different cell types depending on their EMT state.

The final feature we evaluated was cell proliferation. Because it is difficult to assign a growth rate to single cells, we argue that cell proliferation should be measured as the growth rate of the entire cell population. To measure the growth rate of each cell line during the 25-hour period, we employed smoothing splines [41, 42] as shown in figure 4H. Assuming exponential growth in between time points t_*1*_, t_2_, …., t_*n*,_ the growth rate *G*_*CL*_ for each cell line was computed from the fitted spline values according to Equation 1

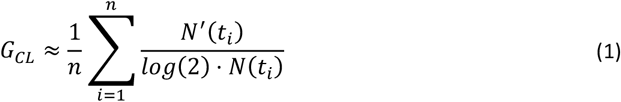

in which *N(t)* = *N*_0_ *exp* (G.t) denotes the number of cells as a function of time and *N*^*′*^(*t*) its derivative. Notably, the doubling time is given by t_2_ = log (2)/G. Our time-course analysis revealed that cells treated with TGF-β1 exhibited the highest growth rate, whereas untreated and EGF-treated MCF7 cells maintained the lowest growth rates across all conditions (Fig. 4I).

### Quantitative assessment of cell plasticity states through multi-parameter analysis

To establish a comprehensive plasticity metric, we analyzed the dataset of individual cells (n=12915) for all tested conditions and calculated their Plasticity Indices (Fig. 5A). The Plasticity Index combines six parameters from morphology, motility and cell growth:

**Figure 5.**
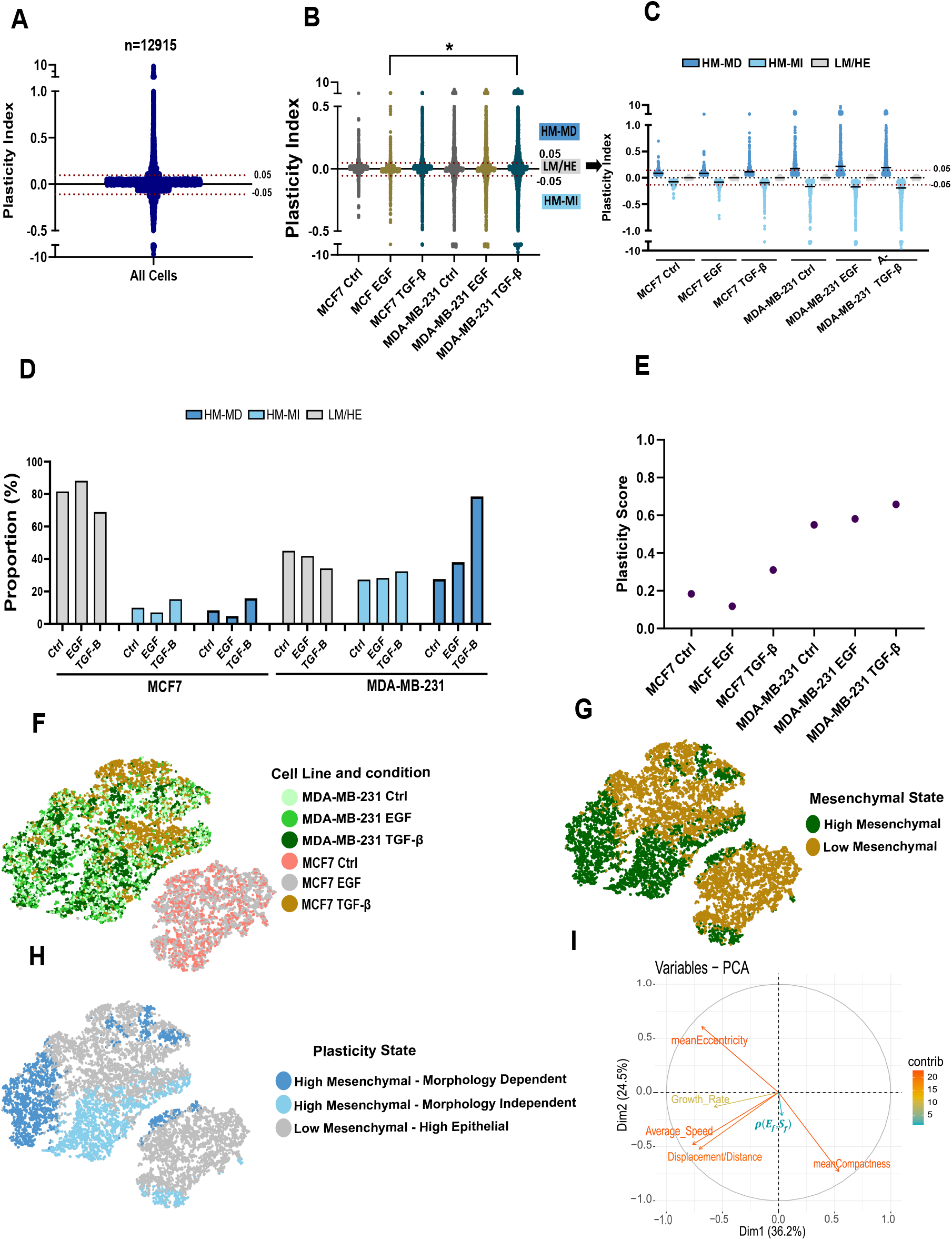
Multi-parameter analysis and classification of cellular plasticity states. **A**) Distribution of plasticity index scores across all analyzed cells (n=12915), showing the range of cellular plasticity in the total population. Horizontal red dotted lines on the y-axis indicate the constant threshold that we use for classification. **B**) Comparative analysis of “Plasticity Indices” across different treatment conditions in MCF7 and MDA-MB-231 cells. **C**) Temporal progression of plasticity parameters across experimental conditions, stratified by plasticity states: HM-MD (High Mesenchymal-Morphology Dependent), HM-MI (High Mesenchymal-Morphology Independent), and LM/HE (Low Mesenchymal-High Epithelial). **D**) Quantification of proportion of cell population for three plasticity states (HM-MD, HM-MI, LM/HE) under different treatment conditions for both MCF7 and MDA-MB-231 cells. Data is presented as a fraction of total cells in each plasticity state. **E**) “Plasticity Score” measured for all conditions of MCF7 and MDA-MB-231. This is the fraction of High Mesenchymal cell categories (HM-MD and HM-MI) in the total cell population of each condition. **F-H**) t-SNE projections based on the combination of six parameters (used to measure the “Plasticity Index” of the cells). Further metadata like Cell line, Mesenchymal state and Plasticity state are projected on t-SNE.I) Principal Component Analysis (PCA) of six parameters of Plasticity index. The plot shows the contribution of each feature in the first two dimensions of PCA. All statistical analyses were performed using Kruskal-Wallis, and data are presented as median where applicable. Significance levels are indicated as follows: ns (not significant): P > 0.05; *: P ≤ 0.05; **: P ≤ 0.01; ***: P ≤ 0.001; ****: P ≤ 0.0001.

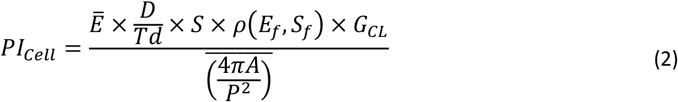

The parameters of Equation 2 include 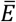 which represents average eccentricity, where eccentricity quantifies how elongated a cell is (it ranges from 0 for a perfect circle to near 1 for an elongated shape). The average compactness represented as 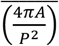 combines perimeter (P) and area (A). The term 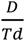 reflects the directionality of cell movement: 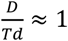 indicates directed, migratory movement observed in mesenchymal cells and 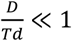 random, undirected movement observed in epithelial cells. *S* indicates average speed of a cell over time, measured in μm/hour. The term *ρ*(*E*_*f*_, *S*_*f*_) denotes the spearman correlation between Eccentricity per frame *E*_*f*_ and Speed per frame *S*_*f*_. Lastly, *G*_*CL*_ denotes the average growth rate (Equation 1).

Using Equation 2, we measured the PI for all 12,915 cells and subsequently classified them within the EMT spectrum. For the classification, we estimated a constant threshold of +/−0.05 for positive and negative PI values, respectively. All cells that have PI > 0.05 were classified as High Mesenchymal-Morphology Dependent (HM-MD), PI < −0.05 indicates High Mesenchymal-Morphology Independent (HM-MI) cells, and all cell with −0.05 < PI < 0.05 were denoted as Low Mesenchymal/High Epithelial (LM/HE). In HM-MD cells, a positive correlation between *E*_*f*_ and *S*_*f*_ indicate that with increasing speed the cells are getting elongated and as the speed decreases the cells become more spherical, hence “Morphology Dependent”. In contrast, HM-MI cells show a negative correlation between *E*_*f*_ and *S*_*f*_ indicating morphology remains independent of speed, justifying the “Morphology Independent” label. Despite this, both HM-MD and HM-MI have high PI values, placing them in the High Mesenchymal class, while LM/HE cells have lower PI values overall.

We further quantified the proportion of cells in each of the three categories—HM-MD, HM-MI, and LM/HE—across both cell line models (Fig. 5D). The proportion of LM/HE cells in MCF7 across all conditions was nearly twice that of LM/HE cells in MDA-MB-231. In contrast, the proportions of high mesenchymal cell categories (HM-MD and HM-MI) were significantly higher in MDA-MB-231 compared to MCF7, a difference that was even more noticeable with TGF-β1 treatment. Additionally, the induction of the mesenchymal state and reduction of the epithelial state were evident in TGF-β1 treated subpopulation groups for both cell lines (Fig. 5D).

Followed by this, we also propose a scoring metric called Plasticity Score (PS_CL_) shown in Equation 3

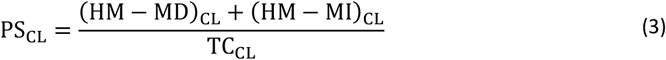

This metric defines the state of cell line model on the EMT spectrum. PS_CL_ close to 1 indicates a cell line model with high mesenchymal cells, thus contributing to strong metastasis potential whereas PS_CL_ close to 0 represents a cell line model with a high content of epithelial cells, therefore low metastasis potential. In Equation 3, the (HM − MD)_CL_ + (HM − MI)_CL_ parameter is the sum of the number of cells from both these categories for the Cell Line (CL) which is then divided by total number of cells TC_CL_ (HM-MD + HM-MI + LM/HE) for the respective cell line. Quantification of overall plasticity scores (Fig. 5E) demonstrated a clear progression from MCF7 control cells (lowest plasticity) to TGF-β1-treated MDA-MB-231 cells (highest plasticity).

t-Distributed Stochastic Neighbor Embedding (t-SNE) algorithm for dimensionality reduction was applied to generate an embedding in two dimensions using the six parameters that were used to compute the PI_Cell_ of all 12,915 cells. Subsequently, we projected meta data of cell lines (Fig. 5F) and the classified mesenchymal states of cells (Fig. 5G-H) on the t-SNE embedding that revealed distinct clustering patterns. The cell line metadata projections shown in Fig 5F is clearly separating the known epithelial cell line MCF7-ctrl and MCF7-EGF from the treated and know mesenchymal cell line conditions like MDA-MB-231. We further checked the contribution of all six parameters by applying PCA and projected the importance of each feature in the first two principal component dimensions (Fig. 5I). The first two dimensions capture about 60% variance in the data. All features contribute to the Plasticity Index that is able to distinguish between the epithelial and mesenchymal cell types.

This analysis provided a quantitative framework for assessing cellular plasticity and demonstrated that while both cell lines possess the capacity for phenotypic plasticity, MDA-MB-231 cells exhibit a greater range and magnitude of plastic responses, compared to MCF7 cells under treatment of TGF-β1.

## Discussion

Our pipeline, Plastimos, offers a novel approach to quantifying EMT-driven cellular plasticity in breast cancer. By integrating live-cell imaging with deep learning-based analysis, we defined numerical Plasticity Indices on a single cell level that capture morphological and behavioral heterogeneity in response to EMT-inducing stimuli. Applying this framework to MCF7 and MDA-MB-231 cells revealed distinct, context-specific plasticity profiles.

We observed differential responses to TGF-β and EGF between the two lines: MCF7 cells upregulated mesenchymal markers under TGF-β, while MDA-MB-231 cells—already mesenchymal—exhibited limited transcriptional changes, likely due to saturation. This aligns with prior findings that highly mesenchymal cells show diminished EMT gene responsiveness[43–45]. Our Plasticity Index uncovered three phenotypic states—High Mesenchymal–Morphology Dependent (HM-MD), High Mesenchymal–Morphology Independent (HM-MI), and Low Mesenchymal/High Epithelial (LM/HE)—which reflect distinct morpho-behavioral characteristics. MDA-MB-231 cells were enriched for HM-MD and HM-MI states, consistent with their inherent plasticity and invasive potential [46, 47].

The relationship between morphology and motility further distinguished the lines: MDA-MB-231 cells showed stronger positive correlations between speed and elongation post-TGF-β treatment, suggesting coordinated morphological adaptation during migration. In contrast, MCF7 cells displayed weaker associations, indicative of limited plasticity. These results are consistent with the studies in the literature, where it is suggested that higher motility in mesenchymal cells is usually associated with a morphological plasticity that is critical for movement through the heterogeneous microenvironments of the tumor [46–49].

Our Plasticity Score effectively quantified cell line–level plasticity, accurately placing MDA-MB-231 cells near 1, with TGF-β treatment yielding the highest score—consistent with our experimental observations. This highlights the potential of Plastimos to reflect ground truth phenotypes. While our six-parameter scoring function captures key morphological and motility traits, incorporating additional features could further enhance its resolution. Given the clinical relevance of EMT-mediated plasticity in therapeutic resistance [50–52], Plastimos may aid in identifying strategies to counteract resistance by targeting EMT dynamics [53–55].

## Acknowledgements

This work was supported by grants from the DFG (HO 6175/4-1, SFB TRR305-A05 and A07 and Deutsche Krebshilfe (70114988). Live-cell imaging data were acquired using the equipment made available by ITEM, Fraunhofer, Regensburg.

## Author’s contribution

S.S. and D.D. conducted the majority of the experiments and analyses. and D.D. and M.H. developed and performed the cell motility analyses. S.M.R and N.S.G performed 3D culture experiments. L.W. assisted with live-imaging experiments. S.S., D.D., and H.H., wrote the manuscript and interpreted the data. H.H. and M.H. supervised the study.

## References

1. Yuan S, Norgard RJ, Stanger BZ. Cellular Plasticity in Cancer. Cancer Discov 2019;9(7):837–51.

2. Cordani M, Dando I, Ambrosini G et al. Signaling, cancer cell plasticity, and intratumor heterogeneity. Cell communication and signaling: CCS 2024;22(1):255.

3. Huang Y, Hong W, Wei X. The molecular mechanisms and therapeutic strategies of EMT in tumor progression and metastasis. J Hematol Oncol 2022;15(1):129.

4. Isert L, Mehta A, Loiudice G et al. An In Vitro Approach to Model EMT in Breast Cancer. Int J Mol Sci 2023;24(9):

5. Buyuk B, Jin S, Ye K. Epithelial-to-Mesenchymal Transition Signaling Pathways Responsible for Breast Cancer Metastasis. Cell Mol Bioeng 2022;15(1):1–13.

6. Bar-Hai N, Ben-Yishay R, Arbili-Yarhi S et al. Modeling epithelial-mesenchymal transition in patient-derived breast cancer organoids. Front Oncol 2024;14:1470379.

7. Huang Z, Zhang Z, Zhou C et al. Epithelial-mesenchymal transition: The history, regulatory mechanism, and cancer therapeutic opportunities. MedComm (2020) 2022;3(2):e144.

8. Anderson RL, Balasas T, Callaghan J et al. A framework for the development of effective anti-metastatic agents. Nat Rev Clin Oncol 2019;16(3):185–204.

9. Jia D, Li X, Bocci F et al. Quantifying Cancer Epithelial-Mesenchymal Plasticity and its Association with Stemness and Immune Response. J Clin Med 2019;8(5):

10. Wang W, Poe D, Yang Y et al. Epithelial-to-mesenchymal transition proceeds through directional destabilization of multidimensional attractor. Elife 2022;11:

11. Bhatia S, Wang P, Toh A et al. New Insights Into the Role of Phenotypic Plasticity and EMT in Driving Cancer Progression. Front Mol Biosci 2020;7:71.

12. Xu D, Wang Y, Wu J et al. Identification and clinical validation of EMT-associated prognostic features based on hepatocellular carcinoma. Cancer Cell Int 2021;21(1):621.

13. Busch EL, Don PK, Chu H et al. Diagnostic accuracy and prediction increment of markers of epithelial-mesenchymal transition to assess cancer cell detachment from primary tumors. BMC Cancer 2018;18(1):82.

14. Meijering E. A bird’s-eye view of deep learning in bioimage analysis. Comput Struct Biotechnol J 2020;18:2312–25.

15. Maška M, Ulman V, Delgado-Rodriguez P et al. The Cell Tracking Challenge: 10 years of objective benchmarking. Nat Methods 2023;20(7):1010–20.

16. Padovani F, Mairhörmann B, Falter-Braun P et al. Segmentation, tracking and cell cycle analysis of live-cell imaging data with Cell-ACDC. BMC Biol 2022;20(1):174.

17. Pylvänäinen JW, Gómez-de-Mariscal E, Henriques R et al. Live-cell imaging in the deep learning era. Curr Opin Cell Biol 2023;85:102271.

18. Nieto MA, Huang RY-J, Jackson RA et al. EMT: 2016. Cell 2016;166(1):21–45.

19. Quinsgaard EMB, Korsnes MS, Korsnes R et al. Single-cell tracking as a tool for studying EMT-phenotypes. Exp Cell Res 2024;437(1):113993.

20. Sheng W, Tang J, Cao R et al. Numb-PRRL promotes TGF-β1 1- and EGF-induced epithelial-to-mesenchymal transition in pancreatic cancer. Cell Death Dis 2022;13(2):173.

21. Liu Q, Sheng W, Dong M et al. Gli1 promotes transforming growth factor-beta1- and epidermal growth factor-induced epithelial to mesenchymal transition in pancreatic cancer cells. Surgery 2015;158(1):211–24.

22. Hao Y, Baker D, Dijke P ten. TGF-β1-Mediated Epithelial-Mesenchymal Transition and Cancer Metastasis. Int J Mol Sci 2019;20(11):

23. Sheng W, Shi X, Lin Y et al. Musashi2 promotes EGF-induced EMT in pancreatic cancer via ZEB1-ERK/MAPK signaling. J Exp Clin Cancer Res 2020;39(1):16.

24. Gottumukkala SB, Ganesan TS, Palanisamy A. Comprehensive molecular interaction map of TGFβ induced epithelial to mesenchymal transition in breast cancer. NPJ Syst Biol Appl 2024;10(1):53.

25. Sachs N, Ligt J de, Kopper O et al. A Living Biobank of Breast Cancer Organoids Captures Disease Heterogeneity. Cell 2018;172(1-2):373-386.e10.

26. Hirokawa Y, Clarke J, Palmieri M et al. Low-viscosity matrix suspension culture enables scalable analysis of patient-derived organoids and tumoroids from the large intestine. Commun Biol 2021;4(1):1067.

27. Livak KJ, Schmittgen TD. Analysis of relative gene expression data using real-time quantitative PCR and the 2(−Delta Delta C(T)) Method. Methods 2001;25(4):402–8.

28. Carpenter AE, Jones TR, Lamprecht MR et al. CellProfiler: image analysis software for identifying and quantifying cell phenotypes. Genome biology 2006;7(10):R100.

29. Pau G, Fuchs F, Sklyar O et al. EBImage--an R package for image processing with applications to cellular phenotypes. Bioinformatics 2010;26(7):979–81.

30. Fares J, Fares MY, Khachfe HH et al. Molecular principles of metastasis: a hallmark of cancer revisited. Signal Transduct Target Ther 2020;5(1):28.

31. Stringer C, Wang T, Michaelos M et al. Cellpose: a generalist algorithm for cellular segmentation. Nat Methods 2021;18(1):100–6.

32. Conner SJ, Guarin JR, L. TT et al. Cell morphology best predicts tumorigenicity and metastasis in vivo across multiple TNBC cell lines of different metastatic potential. Breast Cancer Res 2024;26(1):43.

33. Conner S, Guarin JR, L. TT et al. Cell morphology best predicts tumorigenicity and metastasis in vivo across multiple TNBC cell lines of different metastatic potential. bioRxiv 2023.

34. Liu K, Newbury PA, Glicksberg BS et al. Evaluating cell lines as models for metastatic breast cancer through integrative analysis of genomic data. Nat Commun 2019;10(1):2138.

35. Mandel K, Seidl D, Rades D et al. Characterization of spontaneous and TGF-β1-induced cell motility of primary human normal and neoplastic mammary cells in vitro using novel real-time technology. PLoS One 2013;8(2):e56591.

36. Brünner N, Boysen B, Rømer J et al. The nude mouse as an in vivo model for human breast cancer invasion and metastasis. Breast Cancer Res Treat 1993;24(3):257–64.

37. Jin X, Demere Z, Nair K et al. A metastasis map of human cancer cell lines. Nature 2020;588(7837):331–6.

38. Holliday DL, Speirs V. Choosing the right cell line for breast cancer research. Breast Cancer Res 2011;13(4):215.

39. Katsuno Y, Derynck R. Epithelial plasticity, epithelial-mesenchymal transition, and the TGF-β1 family. Dev Cell 2021;56(6):726–46.

40. Yao W, Wang Z, Ma H et al. Epithelial-mesenchymal plasticity (EMP) in wound healing: Exploring EMT mechanisms, regulatory network, and therapeutic opportunities. Heliyon 2024;10(14):e34269.

41. Everitt B. Book reviews: Chambers JM, Hastie TJ eds 1992: Statisti cal models in S. California: Wadsworth and Brooks/Cole. ISBN 0 534 16765-9. Stat Methods Med Res 1992;1(2):220–1.

42. Green PJ, Silverman BW. Nonparametric Regression and Generalized Linear Models. Chapman and Hall/CRC, 1993.

43. Katsuno Y, Meyer DS, Zhang Z et al. Chronic TGF-β1 exposure drives stabilized EMT, tumor stemness, and cancer drug resistance with vulnerability to bitopic mTOR inhibition. Sci Signal 2019;12(570):

44. Gal A, Sjöblom T, Fedorova L et al. Sustained TGF beta exposure suppresses Smad and non-Smad signalling in mammary epithelial cells, leading to EMT and inhibition of growth arrest and apoptosis. Oncogene 2008;27(9):1218–30.

45. Jiang Y, Woosley AN, Sivalingam N et al. Cathepsin-B-mediated cleavage of Disabled-2 regulates TGF-β1-induced autophagy. Nat Cell Biol 2016;18(8):851–63.

46. Leggett SE, Sim JY, Rubins JE et al. Morphological single cell profiling of the epithelial-mesenchymal transition. Integr Biol (Camb) 2016;8(11):1133–44.

47. Gupta PB, Pastushenko I, Skibinski A et al. Phenotypic Plasticity: Driver of Cancer Initiation, Progression, and Therapy Resistance. Cell Stem Cell 2019;24(1):65–78.

48. Jehanno C, Vulin M, Richina V et al. Phenotypic plasticity during metastatic colonization. Trends Cell Biol 2022;32(10):854–67.

49. Wu J-S, Jiang J, Chen B-J et al. Plasticity of cancer cell invasion: Patterns and mechanisms. Transl Oncol 2021;14(1):100899.

50. Gu Y, Zhang Z, Dijke P ten. Harnessing epithelial-mesenchymal plasticity to boost cancer immunotherapy. Cell Mol Immunol 2023;20(4):318–40.

51. Williams ED, Gao D, Redfern A et al. Controversies around epithelial-mesenchymal plasticity in cancer metastasis. Nat Rev Cancer 2019;19(12):716–32.

52. Santamaría PG, Moreno-Bueno G, Cano A. Contribution of Epithelial Plasticity to Therapy Resistance. J Clin Med 2019;8(5):

53. Hari K, Sabuwala B, Subramani BV et al. Identifying inhibitors of epithelial-mesenchymal plasticity using a network topology-based approach. NPJ Syst Biol Appl 2020;6(1):15.

54. Esquer H, Zhou Q, Nemkov T et al. Isolating and targeting the real-time plasticity and malignant properties of epithelial-mesenchymal transition in cancer. Oncogene 2021;40(16):2884–97.

55. Shi Z-D, Pang K, Wu Z-X et al. Tumor cell plasticity in targeted therapy-induced resistance: mechanisms and new strategies. Signal Transduct Target Ther 2023;8(1):113.

56. Horn LA, Fousek K, Palena C. Tumor Plasticity and Resistance to Immunotherapy. Trends Cancer 2020;6(5):432–41.

## Literature Cited

1. Isert L, Mehta A, Loiudice G, Oliva A, Roidl A, Merkel OM. An In Vitro Approach to Model EMT in Breast Cancer. Int J Mol Sci 2023; 24(9).

