## Supplementary Figures for "Plastimos: a live cell imaging-based framework to study dynamics of EMT-mediated cellular plasticity in breast cancer"

### Figure Supplementary 1

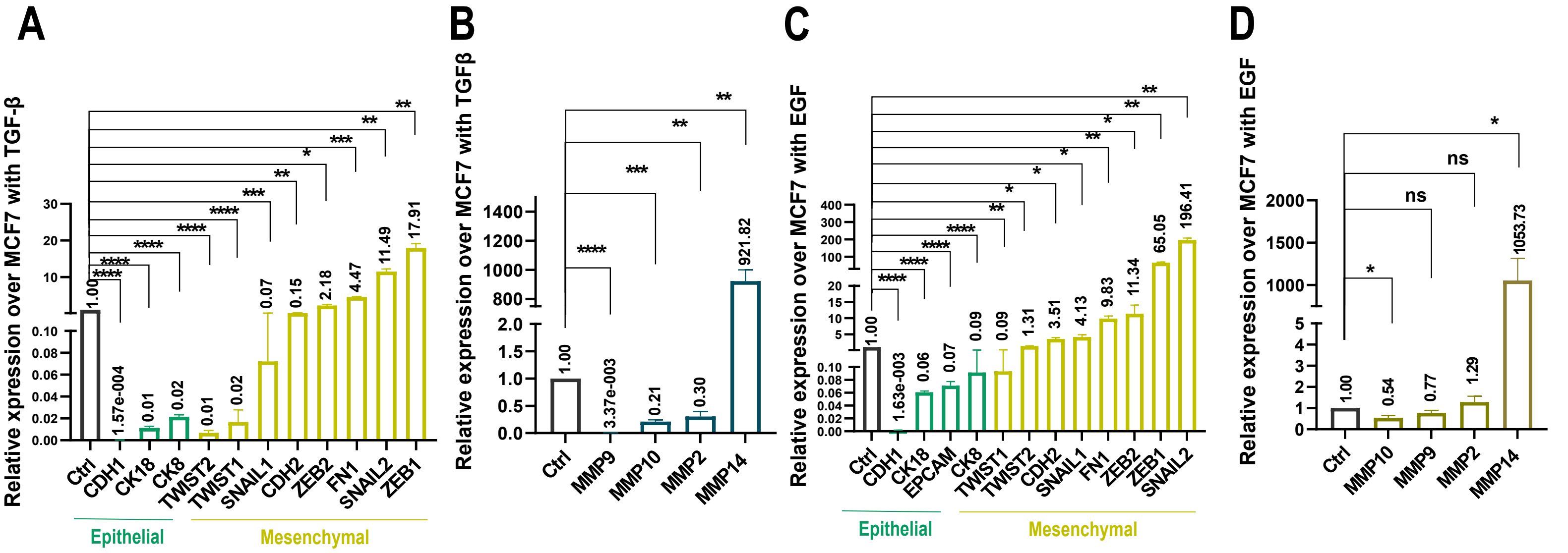


MDA-MB-231 cells + TGF-β (5 ng/mL, 48h)
relative to
MCF7+TGF-β (5 ng/mL, 48h)

MDA-MB-231 cells + TGF-β (5 ng/mL, 48h)
relative to
MCF7 TGF-β (5 ng/mL, 48h)

MDA-MB-231 cells + EGF (100 ng/mL, 48h)
relative to
MCF7+EGF (100 ng/mL, 48h)

MDA-MB-231 cells + EGF (100 ng/mL, 48h)
relative to
MCF7+EGF (100 ng/mL, 48h)

### Figure Supplementary 2

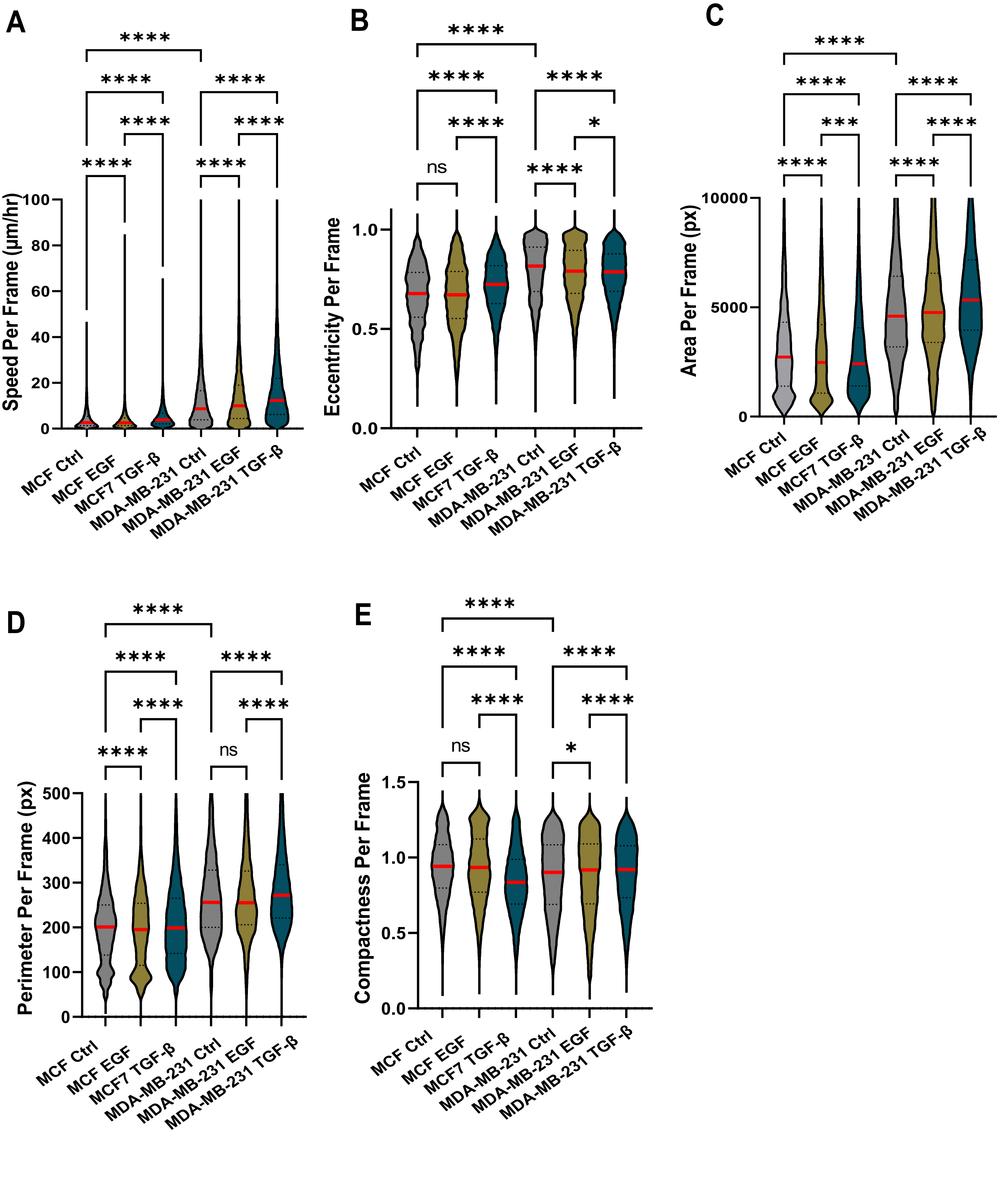

### Figure Supplementary 3

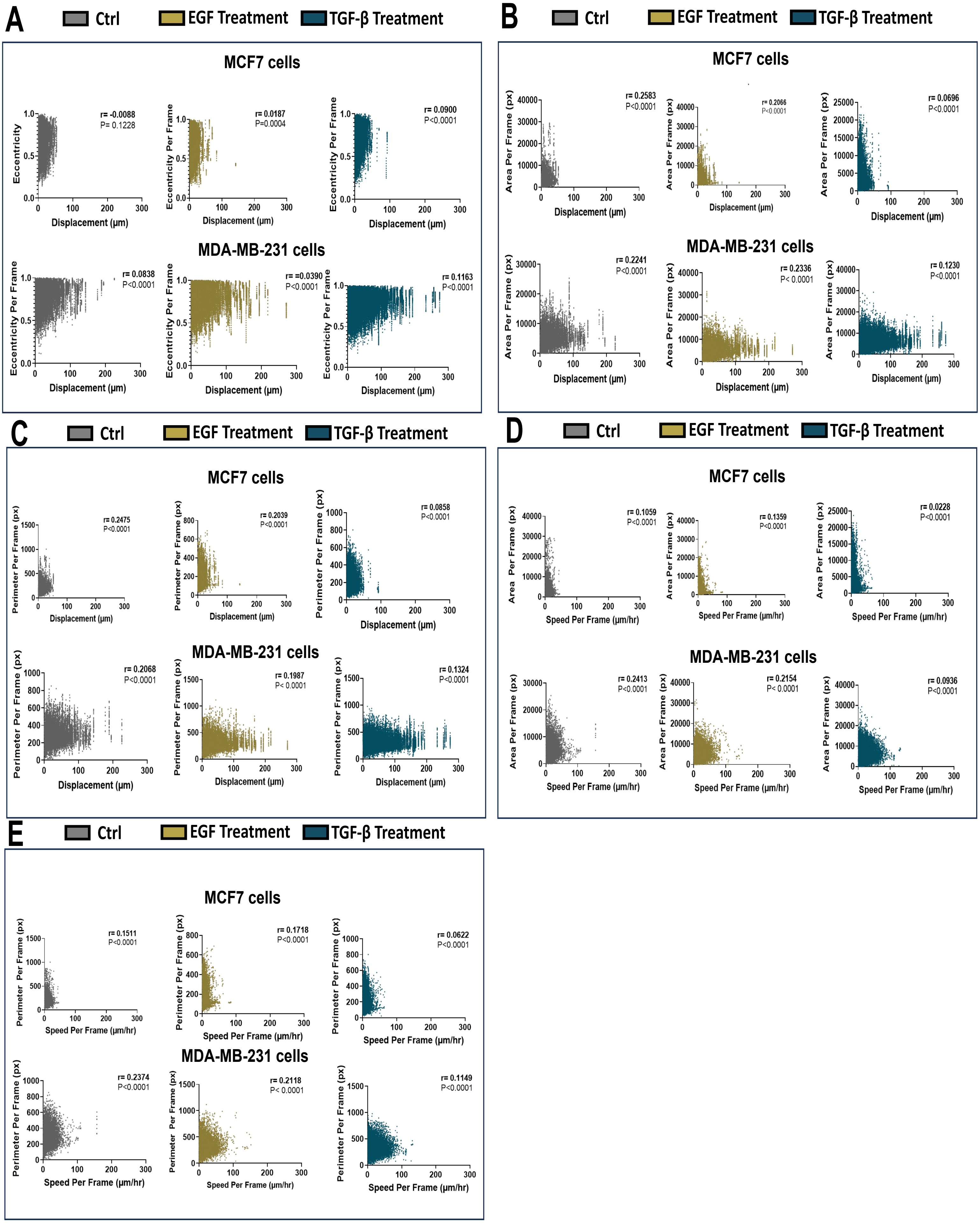
