## Supplementary material for "Plastimos: a live cell imaging-based framework to study dynamics of EMT-mediated cellular plasticity in breast cancer": Legends of Supplementary Figures

**Supplementary Figure 1. EMT induction under TGF-β and EGF treatment. A-B**) Relative quantification (qPCR) of EMT (A) and invasion (B) markers' mRNA expression fold change in MDA-MB-231 cells, treated with TGF-β to MCF7 cells, treated with TGF-β (A). MDA-MB-231 cells have higher expression of mesenchymal markers and lower expression of epithelial markers compared to MCF7 cells. The expression of MMP14 is significantly higher than that of MCF7 cells. **C-D**) Relative quantification (qPCR) of EMT (C) and invasion (D) markers' mRNA expression in MDA-MB-231 cells, treated with EGF to MCF7 cells, treated with EGF. MDA-MB-231 cells have higher expression of mesenchymal markers and lower expression of epithelial markers compared to MCF7 cells. The expression of MMP14 is significantly higher than that of MCF7 cells. Statistical significance in all panels was determined using One Sample t-tests. Significance levels are indicated as follows: ns (not significant): $P > 0.05$; *: $P \leq 0.05$; **: $P \leq 0.01$; ***: $P \leq 0.001$; ****: $P \leq 0.0001$.

**Supplementary Figure 2. Analysis of morphological and motility parameters. A-E**) Analysis of cell speed per frame measured in micrometers per hour (μm/h) (A), Eccentricity per frame (B), Area per frame measured in pixels (px) (C), Perimeter per frame measured in pixels (px) (D) and Compactness per frame (E) across different conditions, demonstrating the velocity differences between cell lines and treatment groups. All data are presented as violin plots showing data distribution, with median values indicated. Comparisons were made between treatment groups and across cell lines using Kruskal-Wallis tests. Significance levels are indicated as follows: ns (not significant): $P > 0.05$; *: $P \leq 0.05$; **: $P \leq 0.01$; ***: $P \leq 0.001$; ****: $P \leq 0.0001$.

**Supplementary Figure 3. Correlation analysis between cell morphology and motility A**) Correlation analysis between eccentricity per frame and displacement of cells in MCF7 (upper panel) and MDA-MB-231 cells (lower panel): control conditions, EGF treatment, and TGF-β treatment. **B**) Correlation analysis between area per frame and displacement of cells in MCF7 (upper panel) and MDA-MB-231 cells (lower panel): control conditions, EGF treatment, and TGF-β treatment. **C**) Correlation analysis between perimeter per frame and displacement of cells in MCF7 (upper panel) and MDA-MB-231 cells (lower panel): control conditions, EGF treatment, and TGF-β treatment. **D**) Correlation analysis between area per frame and speed per frame of cells in MCF7 (upper panel) and MDA-MB-231 cells (lower panel): control conditions, EGF treatment, and TGF-β treatment. **E**) Correlation analysis between perimeter per frame and speed per frame of cells in MCF7 (upper panel) and MDA-MB-231 cells (lower panel): control conditions, EGF treatment, and TGF-β treatment. Gray dots represent control conditions, olive-colored dots represent EGF treatment, and teal-colored dots represent TGF-β treatment. Spearman correlation coefficients (r) and P-values are shown for each condition. All analyses were performed on single-cell measurements across the entire experimental timeframe.
