## Supplementary Table 1 for "Plastimos: a live cell imaging-based framework to study dynamics of EMT-mediated cellular plasticity in breast cancer"

### Table1: Antibodies

| Antibody | Company (Cat-Nr) | Application & Dilution |
|---|---|---|
| Vimentin | Cell Signalling #5741S | 1:100 |
| N-cadherin | Proteintech #22018-1-AP | 1:200 |
| E-Cadherin | Sinobiological #10204-T58 | 1:400 |
| Goat anti-Rabbit IgG (H+L) Highly Cross-Adsorbed Secondary Antibody, Alexa Fluor™ 555 | Invitrogen #A-21429 | IF, 1:200 |
| Rabbit IgG-UNLB | SouthernBiotech #0111-01 | IF, Variable |
