## Supplementary Table 2 for "Plastimos: a live cell imaging-based framework to study dynamics of EMT-mediated cellular plasticity in breast cancer"

### Table 2: Primer Sequences for qPCR

| Name | Sequence For | Sequence Rev | Species |
|---|---|---|---|
| EPCAM | TGTTTGGTGATGAAGGCAGA | CTTCTGACCCCAGCAGTGTT | human |
| CDH1 | TGATTTTTCGGCAGTTCAAGC | ACAATTATCAGCACCCACACA | human |
| CK 18 | GTTCTGCAGATTGACAATGCC | GTCATCAATGACCTTGCGGA | human |
| CK 8 | GATCTCTGAGATGAACCGGAACA | GCTCGGCATCTGCAATGG | human |
| CDH2 | TGGGAATCCGACGAATGG | TGCAGATCGGACCGGATACT | human |
| ZEB1 | GAAAAACCACAAGGGGATGAG | GCTTGACTTTCAGCCCTGTC | human |
| ZEB2 | CCTGCTACTTTCATGCCACC | CCATCAAGCAATTCTCCCTGA | human |
| SNAIL1 | TGCCCTCAAGATGCACATCCGA | GGGACAGGAGAAGGGCTTCTC | human |
| SNAIL2 | ACATGTCGGTTGTCTGGTTG | CGCCAGGAATGTTCAAAGCT | human |
| TWIST1 | CAGCGCACCCAGTCGCTGAA | CTTGTCCGAGGGCAGCGTGG | human |
| TWIST2 | AGCAAGAAGTCGAGCGAAGA | CAGCTTGAGCGTCTGGATCT | human |
| FN1 | CGGTGGCTGTCAGTCAAAG | AAACCTCGGCTTCCTCCATAA | human |
| VIMENTIN | ACTTTGCCGTTGAAGCTGCTA | AAATCTATCTTGCGCTCCTGA | human |
| MMP 14 | GCAGAAGTTTTACGGCTTGCAA | CCTTCGAACATTGGCCTTGAT | human |
| MMP 10 | TCCAGGCTGTATGAAGGAGAGG | GGTAGGCATGAGCCAAACTGTG | human |
| MMP 9 | GGGACGCAGACATCGTCATC | TCGTCATCGTCGAAATGGGC | human |
| MMP 2 | ATAACCTGGATGCCGTCGT | AGGCACCCTTGAAGAAGTAGC | human |
| B-ACTIN | TGGACATCCGCAAAGACCTG | GGGTGTAACGCAACTAAGTCAT | human |
